# A review of Sisyphini Dung Beetles (Coleoptera: Scarabaeinae) endemic to Mauritius: insights from subfossil evidence and phylogenetic analysis

**DOI:** 10.1101/2025.05.16.654365

**Authors:** Federica Losacco, Fernando Lopes, Michele Rossini, Nick Porch, Frank-Thorsten Krell, Saoud Motala, Gimo M. Daniel, Sergei Tarasov

## Abstract

Five indigenous species of Scarabaeinae are known from Mauritius island, classified into two genera: the monotypic *Nesovinsonia* Martínez & Pereira, 1958 and *Nesosisyphus* Vinson, 1946. This study conducts a taxonomic review of the latter genus *Nesosisyphus*, and evaluates its recent extinction using historical and new occurrence records, subfossils, and niche modeling. We provide the first molecular phylogeny and evaluate the evolutionary relationships of *Nesosisyphus* by comparing two different phylogenetic trees: the first, based on a Ultraconserved Elements (UCEs hereafter) dataset, and the second one, on three loci (28S, 16S, COI). We identify five Mauritian *Nesosisyphus* species, including a new extinct one, *Nesosisyphus draco* sp. n., from a 4,000 yr old subfossil. We diagnose, illustrate, and provide an updated key to all the sisyphine species from Mauritius, as well as the occurrence map. Our niche modeling analysis reveals a broader past distribution range for Mauritian sisyphines, now confined to small, disjunct forest patches due to extensive forest contraction and local extinctions.

## INTRODUCTION

The island of Mauritius, together with Rodrigues and La Réunion, forms part of the isolated Mascarene Archipelago in the southwestern Indian Ocean which is globally recognized as one of the most important biodiversity hotspots on Earth (Myers et al., 2000). Approximately 8-10 Mya, Mauritius emerged from a volcanic eruption that remained active until as recently as 25,000 years ago (Montaggioni et al., 1988; Saddul, 1995). Despite covering an area of just ∼ 1800 km^2^, the island’s geological and climatological history, gradually shaped its topography, resulting in a variety of habitat types that support its unique flora (Vaughan and Wiehe, 1937; Strahm, 1996) and fauna (Temple, 1974; Griffiths and Florens, 2006; Motala et al., 2007).

After human colonization in 1598, Mauritius faced extensive habitat destruction (Florens, 2013b), the introduction of alien predators (Bell, 2002), and invasive plants. These human-induced factors led to the extinction of over 130 native species (Florens, 2013a) and posed profound threats to the native ecosystems. By the late 1990s, only about 4.4% of Mauritius retained indigenous vegetation (Hammond et al., 2015), restricted to specific protected areas. However, these remaining forest has suffered considerable degradation and invasion by exotic species (Florens et al., 2017), posing a growing threat to native biota (Monty et al., 2013; Lorence and Sussman, 1986).

Human-induced extinctions in Mauritius have mostly been studied using plants (Virah-Sawmy et al., 2009; Florens et al., 2012) and large vertebrates, such as the iconic Dodo and giant tortoises (Rijsdijk et al., 2011; Cheke and Hume, 2010). However, the impact of these extinctions on insect biodiversity remains poorly understood. Insects play vital roles in terrestrial ecosystems, yet assessing their diversity often requires preliminary taxonomic work. This paper aims to address this gap by conducting a taxonomic review of the endemic roller dung beetles from the tribe Sisyphini, and assessing their recent extinction and diversity through historical and new occurrence records, subfossils, and niche modeling.

Mauritius has two endemic dung beetle genera: the monotypic *Nesovinsonia* Martínez & Pereira, 1958, not yet assigned to any tribe (Tarasov et al., 2016), and *Nesosisyphus* Vinson, 1946, a member of the tribe Sisyphini (Daniel and Davis, 2024). The first sisyphine species discovered from the island was initially classified under the genus *Sisyphus* Latreille, 1807 (Alluaud C., 1898). Later, Jean Vinson discovered three additional species and established a new genus, *Nesosisyphus*, specifically for the Mauritian sisyphines. Consequently, this genus comprised four species: *N. regnardi* (Alluaud, 1898), *N. vicinus* (Vinson, 1939), *N. rotundatus* Vinson, 1946, and *N. pygmaeus* Vinson, 1946.

However, after Vinson’s early research (Vinson, 1939, 1946, 1947, 1951, 1958), there was limited investigation into *Nesosisyphus*. This study, besides Vinson’s materials, is based on two additional data sources: recent dung beetle surveys conducted in 2004 and 2021, which recollected all extant sisyphines except *N. rotundatus*, and a paleontological survey carried out in 2007 in Mare aux Songes (southeast Mauritius) (Rijsdijk et al., 2011). This paleontological survey revealed numerous dung beetle subfossils, including two sisyphines, which are 4200 years old.

While several studies have explored the phylogeny of the tribe Sisyphini (Daniel et al., 2018, 2019; Tarasov and Génier, 2015; Tarasov et al., 2016), none have included the species from Mauritius. Our phylogenetic analysis based on UCEs recovered *Nesosisyphus* as a separate lineage regarding the other groups of sisyphines, while three loci-based phylogeny indicates that *Nesosisyphus* forms a monophyletic group nested within the genus *Sisyphus*. Our taxonomic investigation agrees with Vinson’s previous species concepts, and we did not discover any new extant species on the island. However, the examination of subfossils revealed one new extinct species, *Nesosisyphus draco* sp. n., along with another currently existing species, *N. pygmaeus*. We also identified that *N. rotundatus*, collected in 1940s, may be extinct at present due to habitat loss. These findings indicate that multiple extinction events have influenced the diversity and distribution of *Nesosisyphus* after the human colonization.

Mauritian sisyphines are currently found only in a few isolated patches of mountainous forest throughout the central plateau. We performed niche modeling based on extinct and extant occurrence records to reconstruct their historical distribution and evaluate possible scenarios that may result in such restricted and isolated distribution ranges, which we discuss in more detail below.

This manuscript conforms to the requirements of the amended International Code of Zoological Nomenclature (ICZN) and the new names are not yet registered in ZooBank, the online registration system for the ICZN. Please note that this work is not issued for the purpose of zoological nomenclature and should be considered not published within the meaning of the code (ICZN code, Section 8.2).

## MATERIALS AND METHODS

### Study sites

Data provided in the present paper includes published records from the literature, museum collections, and recent entomological expeditions.

*Nesosisyphus* specimens collected in May-June 2004 and February 2021 come from the Black River Gorges National Park and Moka Mountain Range. Surveys carried out in different areas of the island (eastern side, Montagnes des Créoles and Valleé de l’Est) did not yield any native dung beetles, but only a few specimens of the introduced *Onthophagus unifasciatus*. During fieldtrip in 2021, we used different baits to attract native dung beetles, such as chicken, monkey, tortoise, and human excrement, and carrions (squid and giant snail *Lissachatina fulica*). Additionally, some specimens were obtained through leaf litter sifting and hand collection.

### Morphological investigation and specimen depository

Morphological investigation was primarily carried out on the specimens collected in 2021. Adult morphology was examined on dry-pinned, alcohol-preserved and glycerin-preserved specimens under a Leica S9D stereoscopic microscope (Helsinki) and an Olympus SZX16 (Denver). Specimen preparation in glycerine follows Tarasov and Génier (2015). The whole body of one female and one male specimen of *N. vicinus* and *N. regnardi* were dissected and deposited in MZHF for study and permanent storage.

Examination of the male genitalia involved the following steps: a) dry specimens were softened in a humid chamber prior to extraction of genital capsule; b) clarification in 10% KOH solution for a few minutes and rinse in distilled water; c) dissection of the internal sac using fine forceps; d) examination and photography while immersed in a dense alcohol-based gel; e) genitalia preserved in microvials containing glycerol, each vial pinned with the corresponding specimen.

After morphological examination, all but 21 of the specimens collected in February 2021 were transferred to 96% ethanol for future genetic investigation. The remaining specimens were either pinned or dissected and stored in glycerine, and deposited in the Coleoptera collection of the MZHF.

Examined material is deposited in the following institutes:

DMNS –— Denver Museum of Nature and Science, Denver, Colorado, USA

MNHN –— Muséum National d’Histoire Naturelle, Paris, France

MZHF –— Finnish Museum of Natural History (LUOMUS), Helsinki, Finland

NHML –— The Natural History Museum of London, England

Label data of type specimens are given verbatim; data referring to a single collecting event are separated by commas (“,”) while semicolons (“;”) are used to separate different collecting events.

### Photographs and illustrations

Dorsal habitus of the specimens was photographed with a Canon EOS 5Ds and a Canon MP-E 65 mm macro lens. Male genitalia, body parts of extant species, subfossils and labels were photographed under the Leica S9D stereomicroscope using a Canon EOS M50 Mark II mirrorless camera. Small structures like aedeagi, endophallites and other body parts, were photographed in an alcohol-based hand sanitizer gel used to orientate and fix their position.

All the photographs were stacked with Zerene Stacker 1.04, edited and arranged in plates with Adobe Photoshop 2023. Endophallites were drawn in Adobe Illustrator 2023.

### Archival DNA extraction and sequencing

UCEs sequences from the holotype of *N. rotundatus* were obtained from Lopes et al. (2024). Its DNA was extracted by using a low-cost archival DNA extraction protocol tailored for historical genomic data preserved in insect specimens, and downstream UCE sequencing. Steps of the extraction and methods description can be found in the cited publication.

### Standard DNA extraction and sequencing

We extracted total genomic DNA from: *N. vicinus, N. regnardi, N. pygmaeus*, Sisyphini with Oriental and African distribution. We used the QIAGEN QIAamp DNA Micro Kit following the manufacturer’s protocol. The purified DNA was quantified using Qubit fluorometer 4.0, high-sensitivity reagents, and 3 *µ*l of DNA extract. For the single gene-based phylogeny, we amplified the nuclear gene 28S rDNA domain 2 (28S) and two mitochondrial genes, the 16S rDNA (16S) and the cytochrome *c* oxidase subunit I (COI) using cytiva illustra puReTaq Ready-To-Go PCR, 2 *µ*l of DNA extracts at 20 ng/*µ*l, and 5*µ*l of each primer at 2 *µ*M of concentration. PCR profiles and primers followed Mlambo et al. (2011). PCR products were verified with agarose gel 1.5% and purified using ExoSAP-IT PCR Product Cleanup Reagent (ThermoFisher Scientific), following the manufacturer’s recommendation. Amplicons were sequenced at FIMM Technology Centre (Helsinki, Finland).

Regarding the UCEs-based phylogeny, we used the tailored UCE Scarabaeinae probe-set Scarab 3Kv1 (Gustafson et al., 2023) to produce sequence data for phylogeny reconstruction. See Lopes et al. (2024) for further details on library preparation and sequencing.

### Molecular analysis: UCEs-based phylogeny

We built the UCEs phylogeny by combining our sequences of *N. vicinus, N. regnardi, N. pygmaeus*, African (11 species) and Oriental (4 species) sisyphines, with the dataset containing outgroups from Lopes et al. (2024). We performed the molecular analysis by following the steps described in Lopes et al. (2024).

### Molecular analysis: single gene-based phylogeny

Sequence quality was visually checked with FinchTV 1.5 (Geospiza, Inc., http://www.geospiza.com). Consensus sequences from both strands were generated and manually edited when needed using MegaX (Kumar et al., 2018). Sequenced data were assembled with the Sisyphini dataset containing outgroups (representatives of Epirinini, Coprini, Gymnopleurini, and Eurysternini) from Daniel et al. (2019) using PhylotaR (Bennett et al., 2018), an automated R package (R Core Team, 2021) retrieval of orthologous DNA sequences from GenBank. In addition to the sequenced genes, the nuclear gene carbamoyl-phosphate synthetase 2, aspartate transcarbamylase, and dihydroorotase (CAD) from Daniel et al. (2019) was also added to the final matrix, totalling four gene regions. Sequences were aligned with MAFFT 7.490 (Katoh and Standley, 2013) using the options *–auto* and *–adjustdirectionaccurately*. The best partition and substitution models were selected using ModelFinder (Kalyaanamoorthy et al., 2017) implemented in IQ-TREE 2.2.0.3 (Minh et al., 2020) under Bayesian Information Criterion. Then, a Maximum-Likelihood (ML) phylogeny was recovered with IQ-TREE (Minh et al., 2020), 1000 replicates of ultrafast bootstrap, and Shimodaira–Hasegawa approximate likelihood ratio test (SH-aLRT) (Anisimova et al., 2011). The bootstrapped tree was visualized and edited with FigTree 1.4.4 (Rambaut, 2019). Molecular analysis were carried on for all the *Nesosisyphus* species with the exception of *N. rotundatus*.

### Subfossil sampling

In 2007, a paleontological excavation was conducted in southwestern Mauritius to search for vertebrate remains, like the extinct dodo (*Raphus cucullatus*) and tortoise, to reconstruct the species-rich environment. The fossil site, Mare aux Songes (hereafter MAS), situated in the coastal lowlands (Fig. 1e), was filled with basalt rubble in the mid 20th century as a measure to limit the spread of malaria. Below this rubble the organic sediments preserved in the site are estimated to contain millions of sub-fossil remains (Rijsdijk et al., 2011), including vertebrates, plant macrofossils, and insects. By means of a mechanical digger and hand excavation, three excavation trenches were initially made into the main two MAS basins. Samples of organic sediment were provided to N.P. for insect analysis. Initially three small (<1000 g) samples from the 2007 excavations proved the presence of subfossil insect remains at relatively low densities, but immediately revealed the significance of the material with the occurrence of likely extinct dung beetle taxa, including the species described as new herein. The dating of plant remains and bones from extinct vertebrates, including the dodo and giant tortoises demonstrated that the MAS material dates to the mid-late Holocene, specifically around 4200 kyr BP (Rijsdijk et al., 2011).

**Figure 1.**
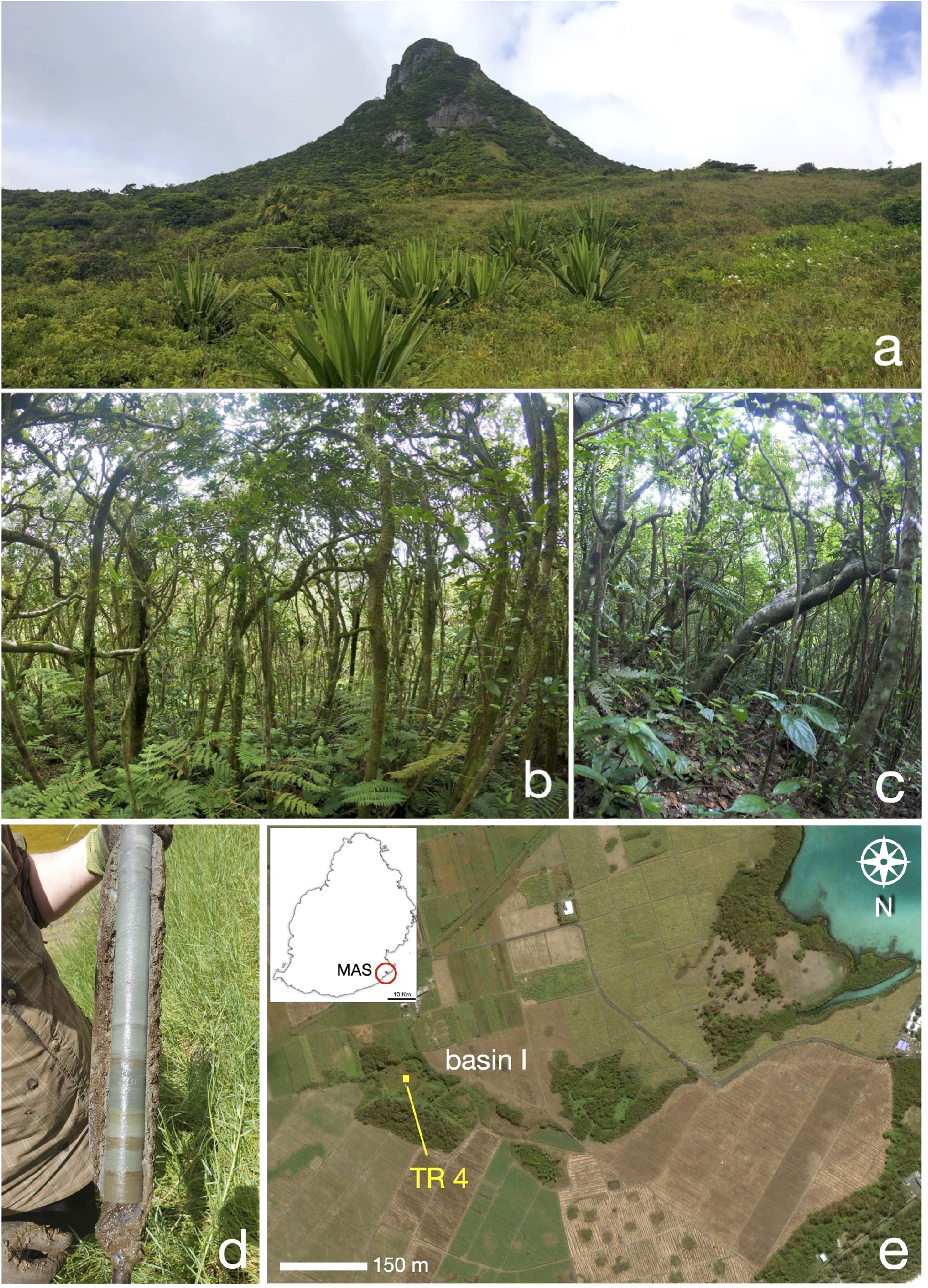
Habitat of *Sisyphus Nesosisyphus* species. (a) View of Mt. Le Pouce, near where *N. regnardi* was collected. (b) Habitat of *N. vicinus*, Mt. Cocotte (SW of Mauritius). (c) Detail of the forest on Mt. Le Pouce, near were *N. regnardi* was collected. (d) Soil excavation from Île d’Ambre (ii.2021) to collect subfossils. (e) Close-up of Mare aux Songes site where the excavation was carried on during the year 2007, modified from Rijsdijk et al. (2009). Subfossils were found in one of the trenches (TR-4) excavated from basin I.

In 2021, 39 additional soil samples were collected by S.T. and M.R. in different Mauritian localities using a Russian corer (Fig. 1d): Kanaka Crater (edge of marsh and forest, 555m asl, 20°24’18.7”S 57°31’05.5”E), Ile d’Ambre (swamp and marsh, 10m asl, 20°02’09.6”S 57°41’49.2”E), Mare la Chaux (dry swamp, 44m asl, 20°12’10.8”S 57°45’10.8”E). Among the several insect remains, only few Aphodiinae subfossil heads and one pronotum were found and none belonging to the subfamily of Scarabaeinae.

### Occurrence data

The occurrence map (Fig. 2) was made in QGIS using the DTM elevation model downloaded from [OpenTopography](https://opentopography.org/) online source and all records from examined material and reliable literature (Vinson, 1939, 1946, 1947, 1951, 1958; Motala and Krell, 2007).

**Figure 2.**
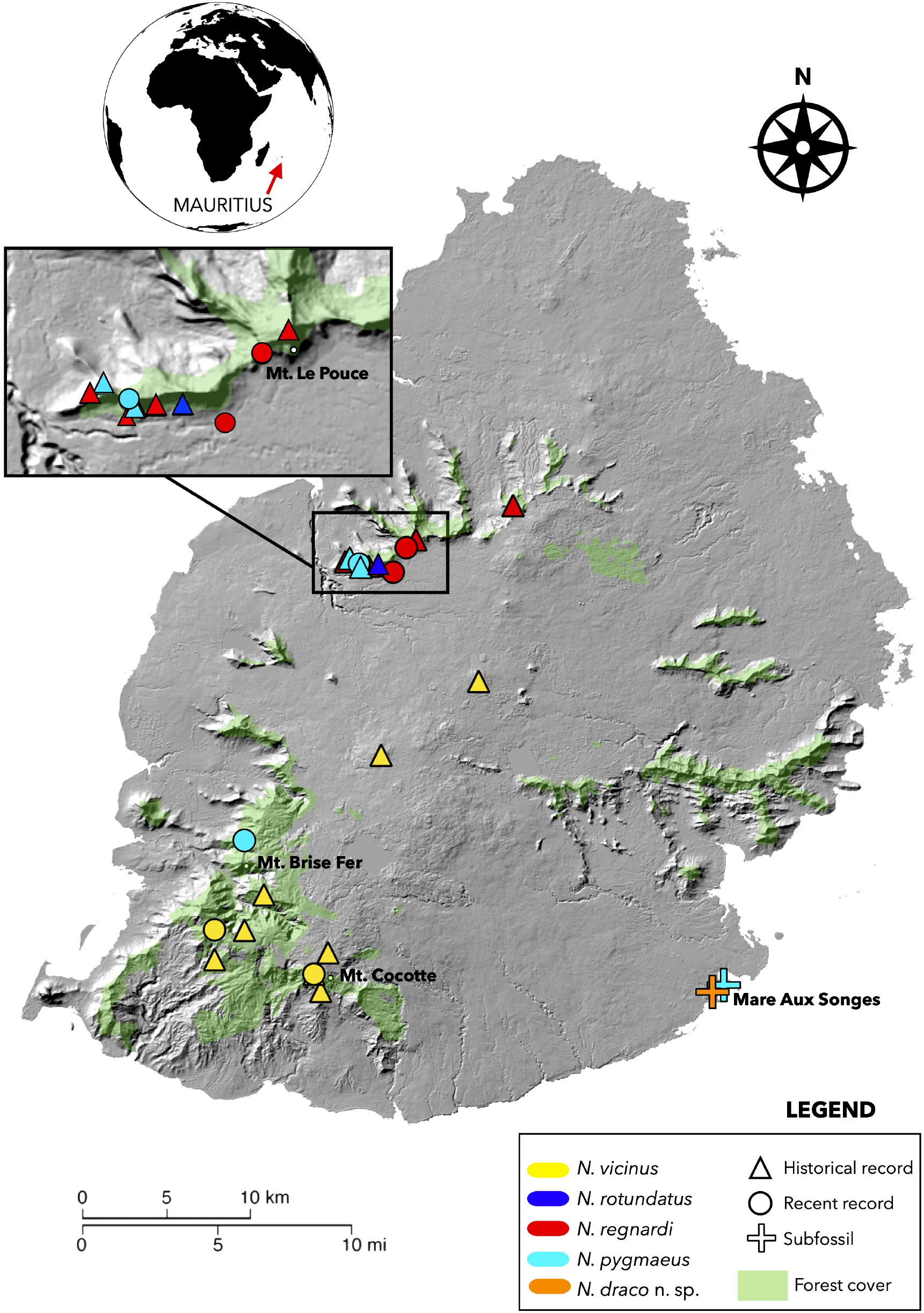
Occurrence map for the Mauritian species of *Nesosisyphus*. Records subdivided per historical (pre-2000s), recent (post-2000s) and subfossil data. Forest cover (from 1997 map) modified from Buckland et al. (2014)

**Figure 3.**
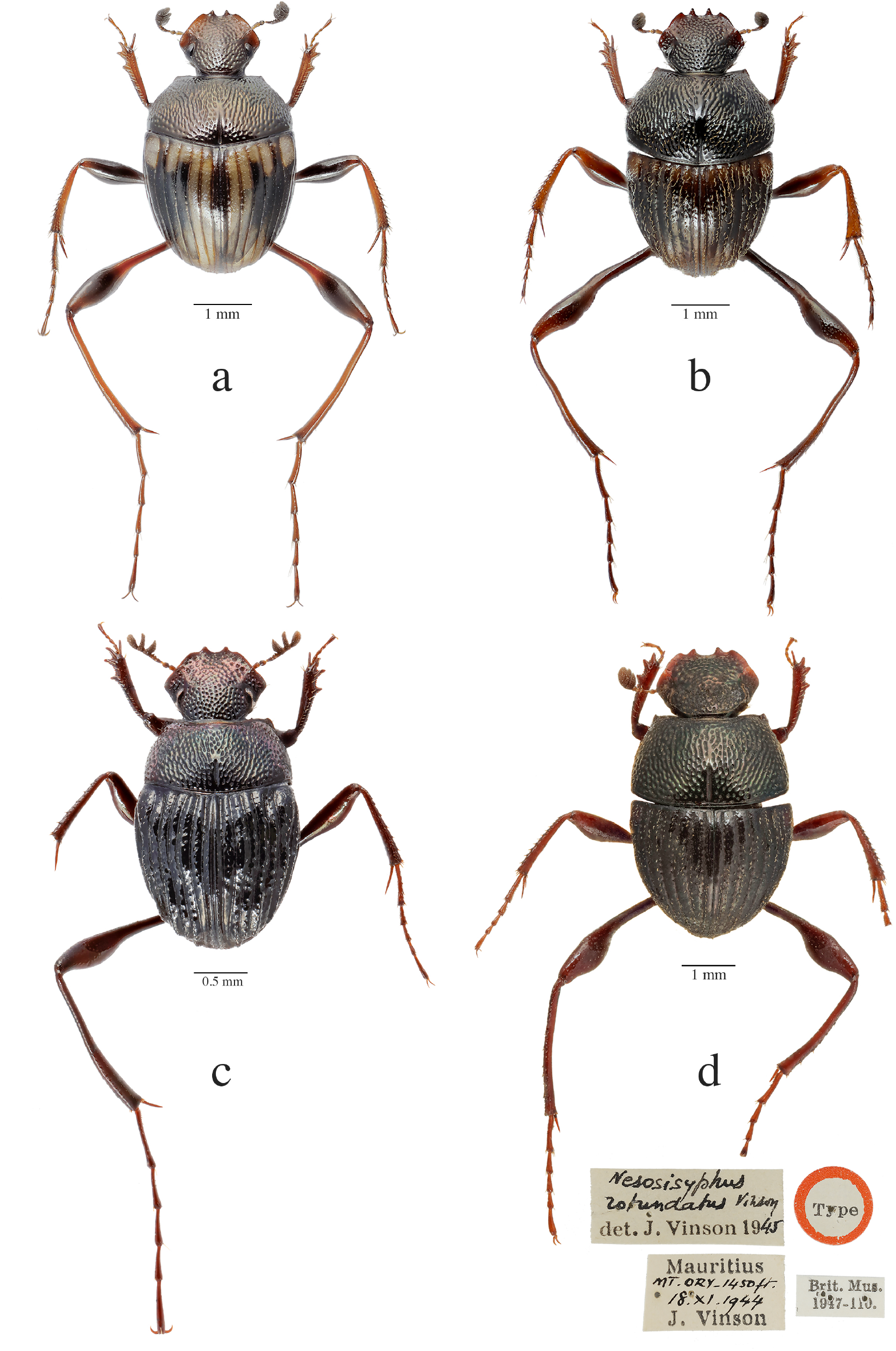
Habiti of extant *Nesosisyphus* species. (a) *N. vicinus* (Vinson, 1939); (b) *N. regnardi* (Alluaud, 1898); (c) *N. pygmaeus* Vinson, 1946; (d) holotype of *N. rotundatus* Vinson, 1946, with original labels.

### Niche modeling

We used historical and recent occurrences to assemble a dataset for all Mauritian sisyphines. Altogether, we had 29 records of presence corresponding to 25 different spatial coordinates. Because of the small number of observations we ran niche modeling only for (1) *N. vicinus*, (2) *N. regnardi*, (3) all extant *Nesosisyphus*, and (4) all extant *Nesosisyphus* and the subfossils. We downloaded bio-climatic layers for the period 1970-2000 at a 30s (1*km*^2^) resolution from the WorldClim database (Fick and Hijmans, 2017). Niche modeling was done using the biomod2 R package (Thuiller et al., 2023). We used presence only data and therefore simulated pseudo-absence data. To have a full coverage of the island, and given we had at most 25 observations of presence only, we ran two sets of simulations with 100 pseudo-absence data points, and with two sets of 50 points. Given the low amount of data available we restricted the niche modeling to only two climatic layers. We retained the annual mean temperature (BIO1) and the annual precipitation (BIO12). We used 11 different algorithm: ANN, CTA, FDA, GAM, GBM, GLM, MARS, MAXNET, RF, SRE, XGBOOST, with the default parameters. Alos, we used 80% of our data for calibration, hence, keeping 20% for model validation. Ensemble models were created using ‘EMmean’, ‘EMcv’, ‘EMci’, ‘EMmedian’, ‘EMca’, ‘EMwmean’.

### Baiting experiment

Feeding preferences for *N. pygmaeus* were assessed in the upland of Brise Fer forest (SW Mauritius) using 8 types of baits (chicken dung, bat dung, monkey dung, pig dung, crocodile dung, tortoise dung, rotting chicken, rotting snail) in both managed and adjacent non-managed plot, during the year 2004 (Motala, 2004). The baits were suspended using meshwire net over the pitfall traps and left to run for 24 hours before being replaced with fresh baits. Traps were spaced by 3 meters and were implemented with raincovers. The traps were devoid of preservatives in order to avoid interference with the population size. Beetles in the trap were counted before being released 10 m from traps to minimise the interference of trap-addiction. Seven replicates of the baiting regime were carried out.

The count data were analysed using the non-parametric Friedman’s test with SsS version 1.0 (Rubisoft Software GmbH). We compared the totals in managed and unmanaged forests using an exact permutation test (Randomization Test U in SsS). We treated these totals as independent pooled samples.

## RESULTS

### Phylogenetic relationships

We obtained all the data available from Museum collections and previous entomological expeditions, as well as fresh material for molecular analysis. We extracted DNA from: *Nesosisyphus vicinus, N. regnardi, N. pygmaeus*, Sisyphini with Oriental (4 species) and African (13 species: *S. sordidus, S. muricatus, S. umbraphilus, S. australis, S. manni, S. barbarossa, S. spinipes, S. infuscatus, S. calcaratus, S. rubrus, S. mirabilis, S. gazanus, S. inconspicuus*) distribution.

#### UCEs-based phylogeny

This phylogeny is complete of all the species of *Nesosisyphus*, including *N. rotundatus*, and the 4 species native to Oriental region (Laos), (Fig. 4). The tribe Sisyphini is recovered as a sister clade to Epirinini tribe, and it splits into two different groups: clade A (BS = 100%) is exclusive to all the *Nesosisyphus* species, and sister group to the clade B, that comprises all the remaining lineages of Sisyphines.

**Figure 4.**
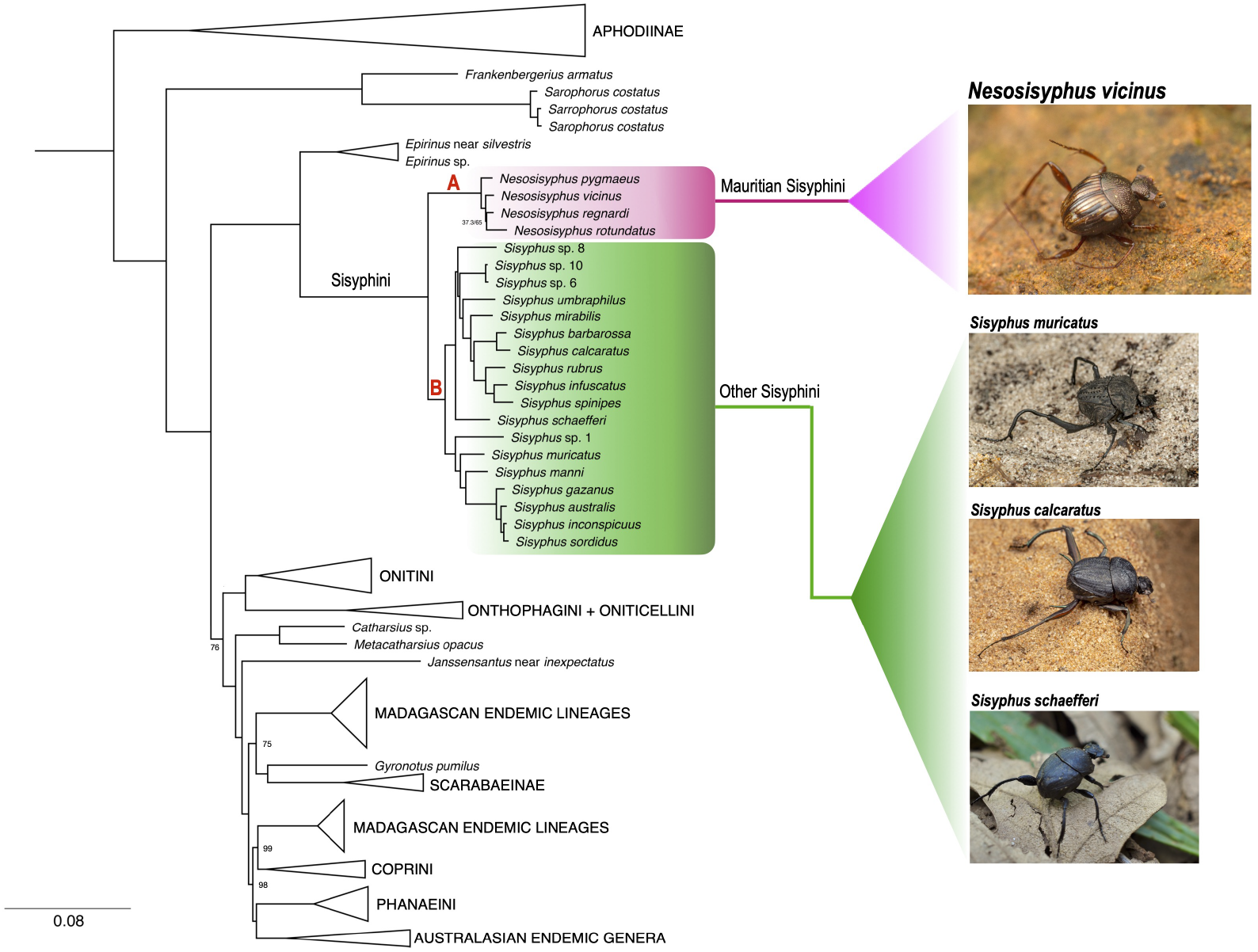
Phylogenetic tree using UCEs shows the systematic position of *Nesosisyphus* (Clade A, highlighted in purple). Other sisyphines are from Oriental, African and Palearctic region (Clade B, highlighted in green). Bootstrap values are indicated when minor to 100; all other nodes have full support.

#### Single gene-based phylogeny

Our ML phylogeny recovers the tribe Sisyphini as monophyletic with bootstrap support of 100% (Fig. 5). It splits into two clades. Clade A (BS = 97%) includes two subgenera which were recovered monophyletic: *Neosisyphus* (BS = 85%) and *Sisyphus* (BS = 53%). All the species in clade A occur in the Afrotropics, except *Sisyphus schaefferi*, which is distributed in the Palearctic.

**Figure 5.**
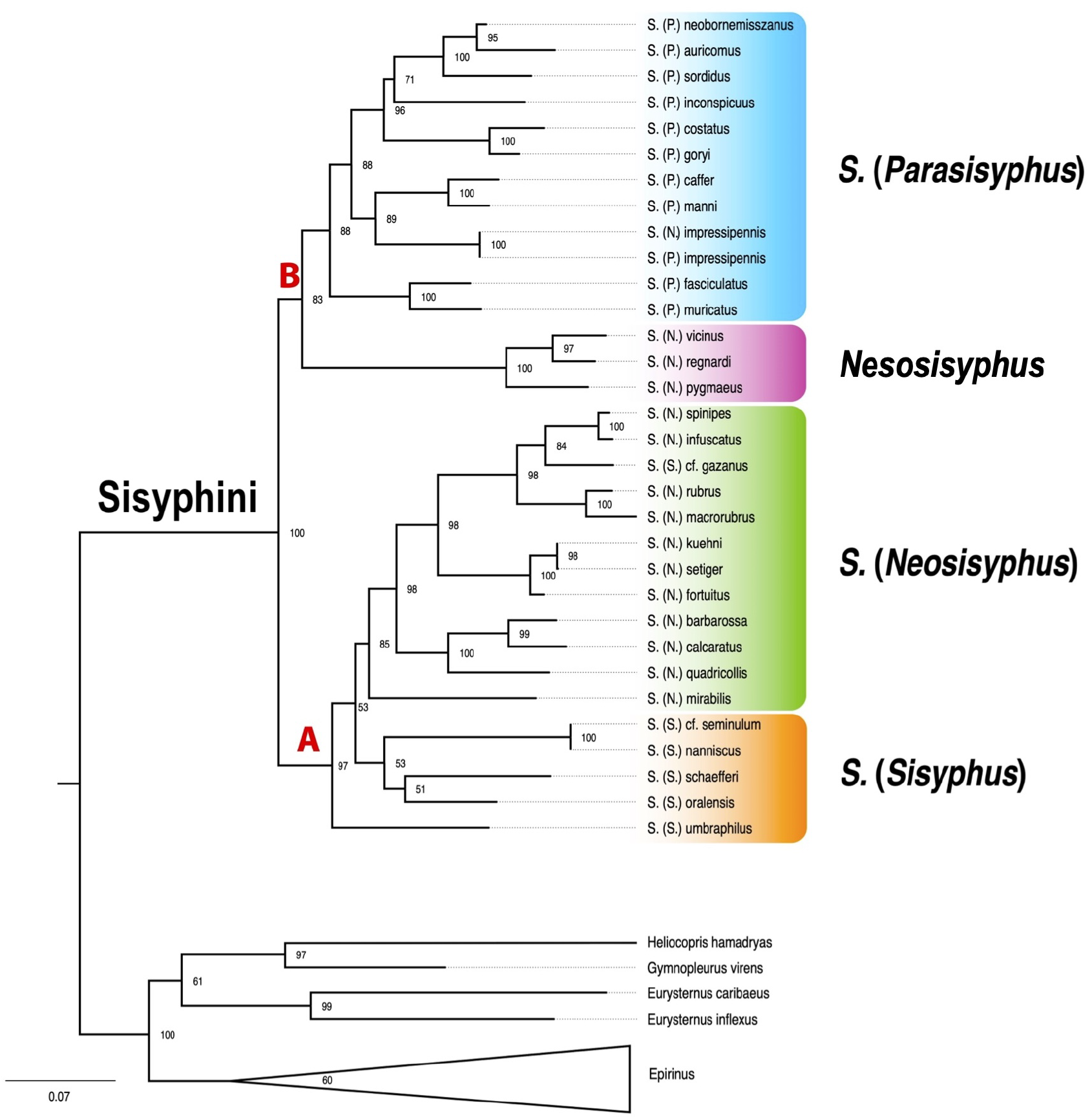
Concatenated phylogenetic tree using three loci (28S, 16S, COI) shows the systematic position of the Mauritian sisyphines (highlighted in purple) nested within the Sisyphini clade and sister group of *S*. (*Parasisyphus*) (Clade B). Maximum likelihood bootstrap are presented for *Sisyphus* lineages and outgroups.

Clade B (BS = 83%) consists of two subclades: the Afrotropical subgenus *Parasisyphus* (BS = 88%) and the Mauritian *Nesosisyphus* (BS = 100%). While *Nesosisyphus* is well-supported as monophyletic, it is nested within the genus *Sisyphus*, which includes African and Palaearctic species in this study.

We report some possible misidentifications in GenBank data that should be taken into account when interpreting the phylogeny and the evolutionary relationships of Sisyphini: only female specimens of *S*. (*N*.) *kuehni* and *S*. (*N*.) *setiger* were examined (Daniel et al., 2020, 2019), which may have led to misidentifications as shown by a lack of genetic differences in the tree; the sequences of *S. seminulum* and *S. nanniscus* in GenBank are from South Africa, but *S. seminulum* does not occur in this region, suggesting a possible misidentification or mislabeling, especially considering that *S. nanniscus* was previously placed as junior subjective synonym of *S. seminulum*, until recently when *S. nanniscus* was revalidated (Montreuil, 2017; Daniel et al., 2018); similarly, the sequence of *S. gazanus* was deposited in GenBank with a possible misidentification. The species is not typically found in Southern Africa, (Monaghan et al., 2007), but is more commonly associated with tropical regions such as Zimbabwe, Malawi, and Mozambique (Daniel et al., 2018).

### Niche modeling

We considered both the ensemble models, which combine all models based on their TSS and ROC scores, and the SRE model. The SRE model provides broad ecological niches by selecting areas at the intersection of the minimum and maximum observed values for each layer used in niche modeling (Fig. 6B). We chose this approach to account for the limited number of data points in our dataset.

**Figure 6.**
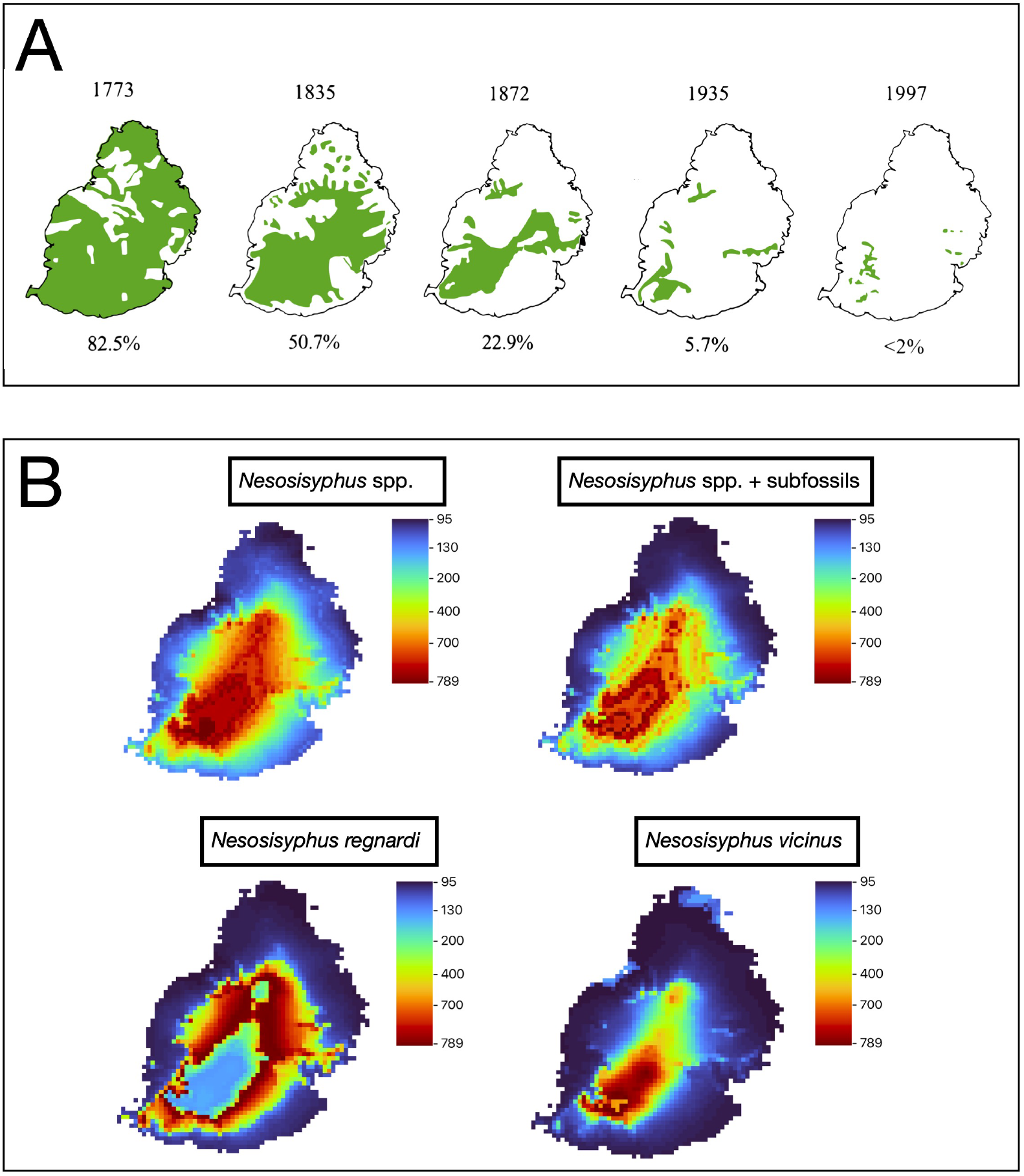
(A) Changes in the native forest cover in Mauritius island (green areas indicate places with > 50% of native plant species). Modified from Cheke and Hume (2010). (B) Ecological niche model (Surface Range Envelope, SRE) estimated suitable areas for *Nesosisyphus* species with and without subfossil (top row), and of *N. regnardi* and *N. vicinus* (bottom row). Red areas indicate a high probability of species presence, while blue areas show a less probability of species occurrence.

Our analysis of *N. vicinus* and *N. regnardi* alone showed that these species inhabit distinct habitats in the central plateau, with very little overlap in their distribution ranges. When we modeled the *Nesosisyphus* dataset, which includes all extant species, we found that its potential niche is widespread across the central plateau, and does not extend to the lowland periphery. Interestingly, adding subfossil data from lowland area did not significantly alter the predicted distribution. It is worth noting that the distribution slightly contracted, likely due to the subfossil data being an outlier.

### Feeding preferences

From the baiting experiment, the dung preference for *Nesosisyphus pygmaeus* could not be detected statistically (due to small number of trapped specimens). The null hypothesis, stating that beetles show no preference for any particular bait, was rejected. This conclusion is based on the results of Friedman’s test, which identified a significant difference in the count data (Fr = 12.179; Frcorr = 17.381; p = 0.022). However, the test did not specify which combinations of baits were different. Notably, tortoise dung was the only bait that did not attract any dung beetles. The count data can be found in supplementary material.

## TAXONOMY

### Updated identification key to the adults of *Nesosisyphus* species, modified from Vinson (1946)

1. Edge between clypeal teeth with a small protrusion (Fig. 7f); species known only by one subfossil head collected in a fossil bed in Mare aux Songes (southeast coast of Mauritius)… … … … … … … … … … … … … … … … … … … … … … … … … … . . ***Nesosisyphus draco*** n. sp.

– Edge between clypeal teeth straight or curved, without protrusion. Extant species. **2**

2 Clypeus bidentate, margin between the teeth semi-circular; metalegs with tarsus longer than tibia; hind wings fully developed **3**

– Clypeus quadridentate, margin between medial teeth straight; metalegs with tarsus shorter or equal to tibia; hind wings brachypterous **4**

3 Elytra medium brown to pale yellow, coloration as in Fig. 8a; surface of elytra smooth and glabrous with isodiametric microsculpture; male genitalia as in Fig. 9a; occurs in the forested slopes of the south-western plateau; size relatively larger: 3.4 to 4.4 mm… … … … … … … … … … … … … … … … … … … … … … … … … … … … … . . ***Nesosisyphus vicinus*** (Vinson, 1939)

– Elytra uniformly black, moderately shining; surface appears irregular and lumpy, without microsculpture; interstriae strongly convex, almost costulate; humeral and apical calli prominent, the latter protruding backward in a small tubercle; male genitalia as in Fig. 9d; occurrence records from Mt. Le Pouce and Brise Fer forest; size smaller: 2.3 to 3.2 mm … … … … … … … … … … … … … … … … … … … … … … … … … … . ***Nesosisyphus pygmaeus*** Vinson, 1946

4 Elytra with distinct humeral and apical calli; microsculpture with isodiametric meshes gently impressed; hind tarsus equals the tibia in length; hind wings reduced to three-fourths of elytra; male genitalia as in Fig. 9b; species found across the Moka mountain range in the north-west of the island: Mt. Le Pouce, Mt. Ory, Mt. Calebasses; size: 3.3 to 4.9 mm … … … … … … … … … … … … … … … … … … … … … … … … … … . ***Nesosisyphus regnardi*** (Alluaud, 1898)

– Elytra without calli or depressions; microsculpture absent; hind tarsus shorter than tibia; hind wings reduced to one-third of elytra; male genitalia as in Fig. 9c; species known only by the type series collected on the S slope of Mt. Ory; size: 3.5-3.9 mm … … … … … … … … … … … … … … … … … … … … … … … … … … … … … … . ***Nesosisyphus rotundatus*** Vinson, 1946

**Figure 7.**
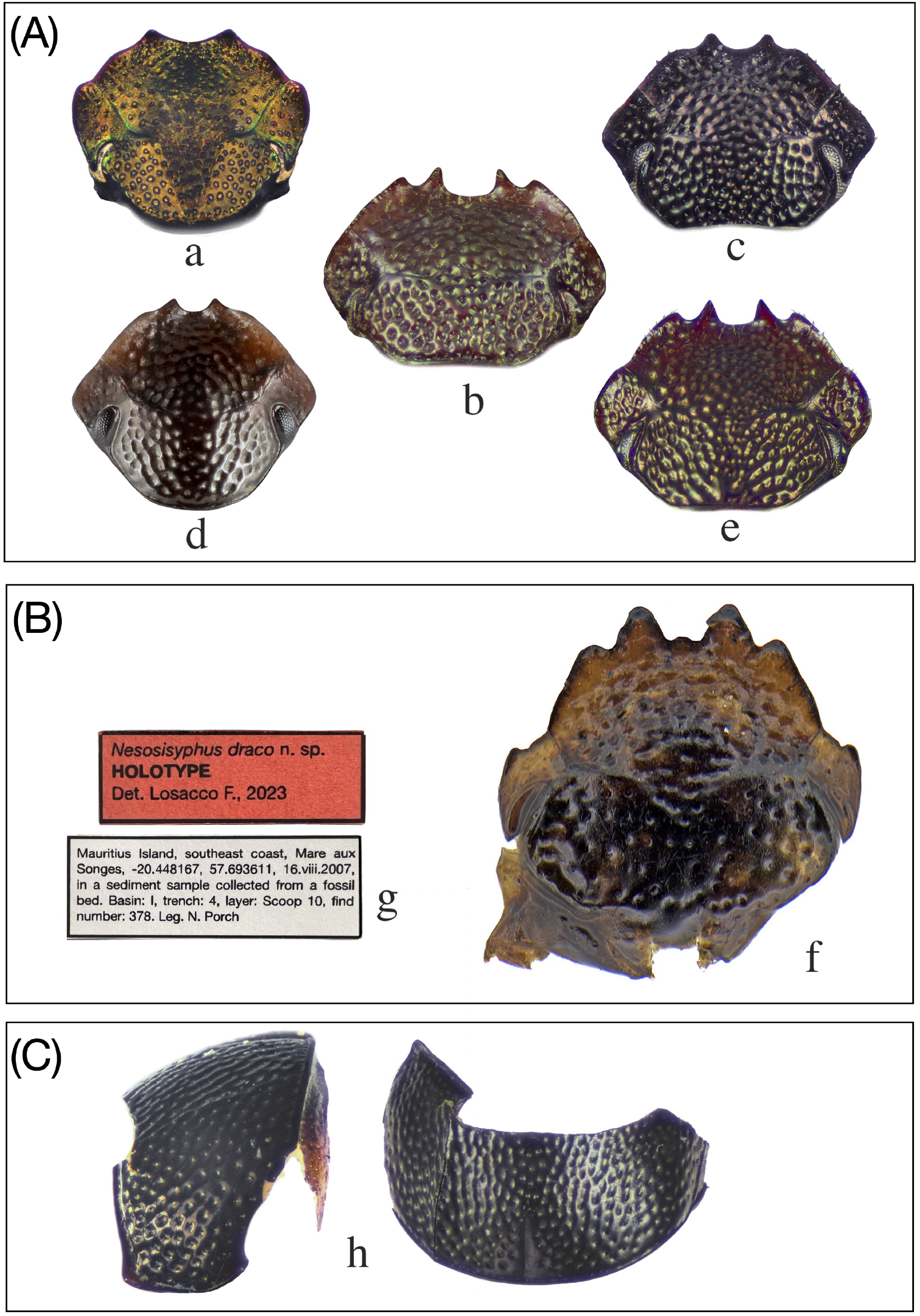
(a-f) Morphological differences in heads between *Nesosisyphus* and *S*. (*Sisyphus*) species. (A) Extant species: (a) *S*. (*S*.) cf. *neglectus* from Oriental region (Laos); (b) *N. rotundatus*; (c) *N. pygmaeus*; (d) *N. vicinus*; (e) *N. regnardi*. (B) New subfossils species from Mare aux Songes (SE Mauritius): (f) *N. draco* n. sp., holotype and type labels (g). (C) Subfossil pronotum (lateral and dorsal view) from Mare aux Songes (SE Mauritius): (h) *N. pygmaeus*.

**Figure 8.**
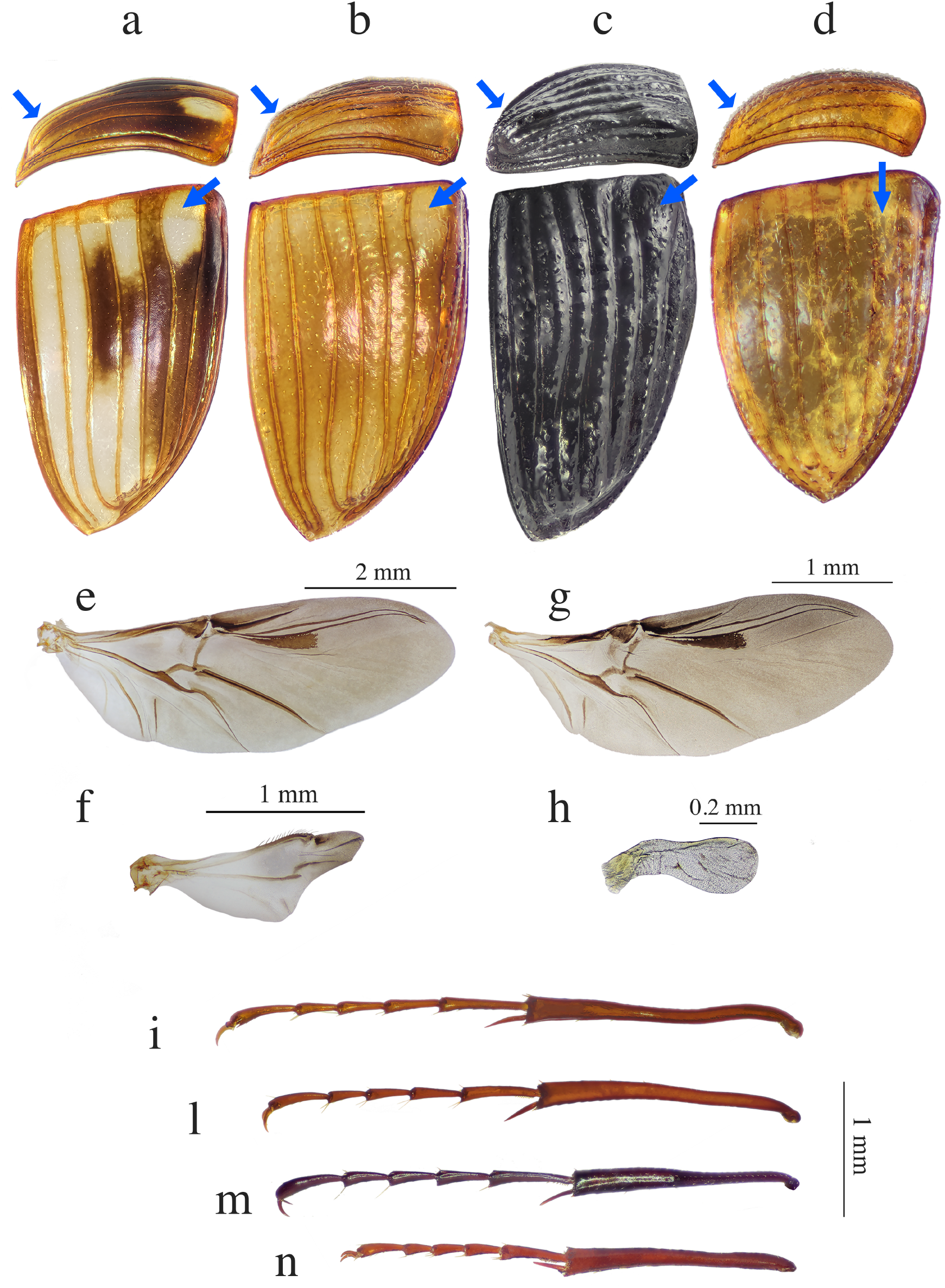
Morphological differences in body parts between *Nesosisyphus* extant spp. For each species, in alphabetical order, are illustrated: right elytra (lateral and dorsal view) with blue arrows pointing at humeral and apical calli which are absent in (d); right hind wing; left hind tarsus and tibia (ventral view). *N. vicinus* (a, e, i); *N. regnardi* (b, f, l); *N. pygmaeus* (c, g, m); *N. rotundatus* (d, h, n).

**Figure 9.**
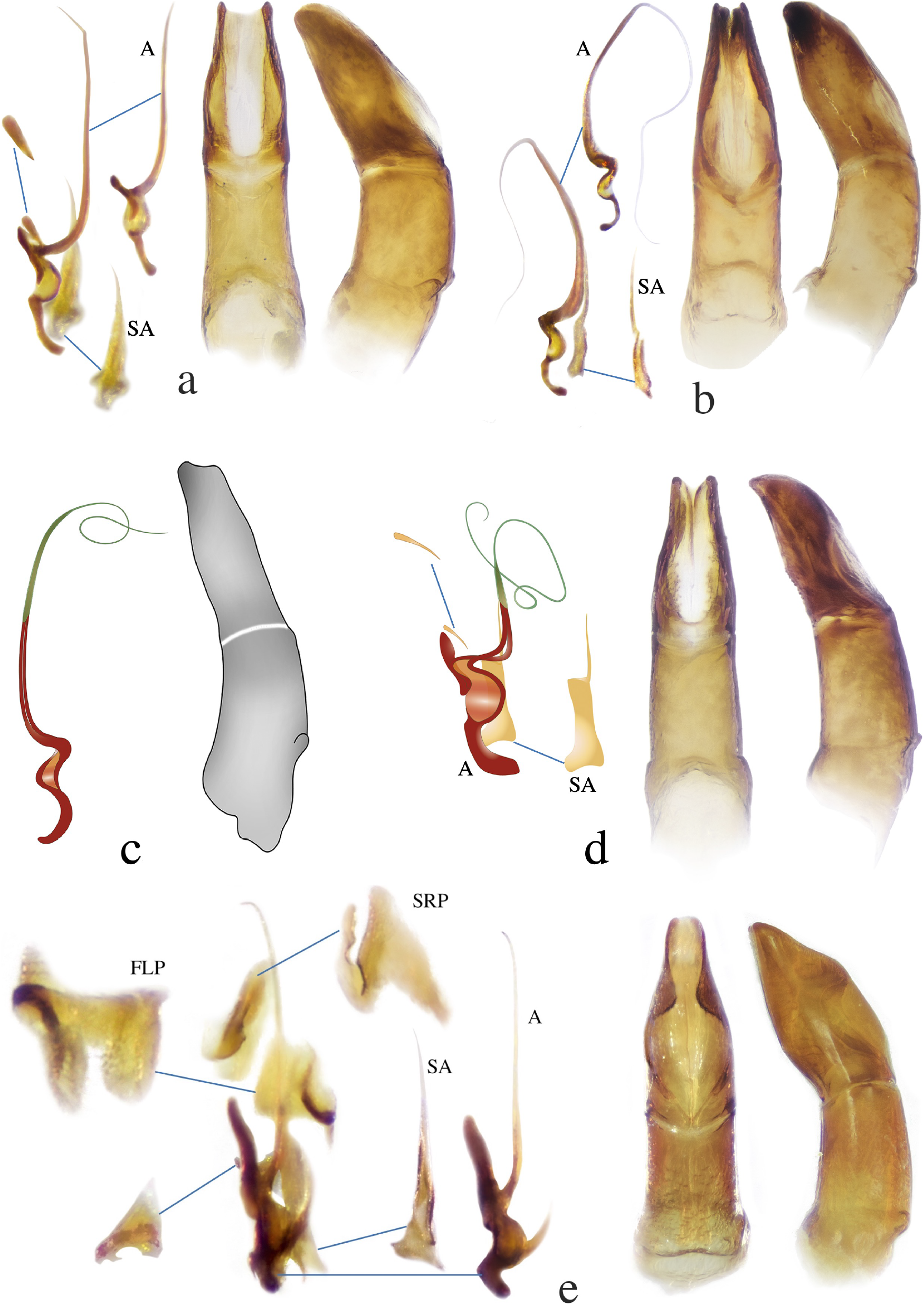
Morphological differences in male genitalia between *Nesosisyphus* and *S*. (*Sisyphus*) species. For each species are shown: parameres (dorsal view), aedeagus (lateral view), and endophallites (acronyms follow Tarasov and Génier (2015)). (a) *N. vicinus*; (b) *N. regnardi*; (c) *N. rotundatus*, modified from Vinson, 1946; (d) *N. pygmaeus*; (e) *S*. (*S*.) cf. *neglectus* from Oriental region (Laos).

### *Nesosisyphus* Vinson, 1946

*Nesosisyphus*: Vinson (1946): 90

*Nesosisyphus*: Vinson (1951): 93

*Nesosisyphus regnardi*: Edmonds and Halffter (1978): 325, 328

#### Type species

*Sisyphus vicinus* Vinson, 1939, by original designation

#### Diagnosis

The genus *Nesosisyphus* is morphologically similar to Sisyphus (Sisyphus), however, we can distinguish them as follows: (i) pronotal disc not concave laterally in *Nesosisyphus*, it bears only a well-marked line along its lateral region; (ii) in *Nesosisyphus* the pronotum is deeply impressed by oblique punctures that converge towards the center of its base, while in *Sisyphus* they are ocellate or simple; (iii) elytra convex with lateral margin slanting downwards in *Nesosisyphus* and not upward towards the apex like in *Sisyphus*; (iv) average body size smaller (between 2.3 and 4.9 mm) in *Nesosisyphus*; (v) endophallites without the fronto-lateral peripheral sclerite (FLP) and the superior right peripheral sclerite (SRP) in all the *Nesosisyphus* species, as illustrated in Fig. 9.

#### Distribution and ecology

*Nesosisyphus* is endemic from Mauritius island. Extant species of *Nesosisyphus* are distributed on the montane tropical forest in the western side of Mauritius island. Present-day populations are living in climate conditions that characterize the uplands habitats, such as lower temperatures and high humidity. Reproductive behaviour remains unknown. Specimens can be collected using dung-baited pitfall traps, leaf litter sifting, and hand collection.

### *Nesosisyphus draco* n. sp

Zoobank link:

Figs. 7f-g

#### Type locality

Mare aux Songes, Mauritius

#### Size

Head: 0.7 mm.

#### Examined type material

(holotype, MZHF) Mauritius Island, southeast coast, Mare aux Songes, -20.448167, 57.693611, 16.viii.2007, in a sediment sample collected from a fossil bed. Basin: I, trench: 4, layer: Scoop 10, find number: 378. Leg. N. Porch.

#### Diagnosis

Clypeus with two pairs of teeth, medial and lateral. A small protrusion is separating the medial teeth. The size of the head is similar to *N. pygmaeus*, but its shape is different due to the extra pair of teeth.

#### Description

Head, oval-shaped, narrowing from base to apex, with two rounded swells on the vertex; covered with scattered deep punctures, some very shallow and barely visible punctures on lateral lobes; lateral lobes narrowing onward with sharp angles on apical half; clypeus not upcurved, bearing four triangular, apically quite blunt teeth, small protrusion separating medial teeth which are more prominent than lateral ones; eyes reduced, almost indistinct on the upperside.

#### Distribution and horizon

The specimen was found in basin I in MAS (see Fig. 1e). The sediments in MAS were from a shallow lake/marsh supplied by groundwater, not surface water from rivers or streams. This suggests that the subfossil is local, likely falling into the site and getting preserved. The site dates to 4200 kyr BP. It appears this species lived in lowland areas and, possibly, went extinct between the 16th and 18th centuries due to the loss of lowland habitat.

#### Remarks

Although we only found one body part of this species among the subfossils from MAS, its distinct morphological features leave no doubt that it is a new species. The head is well-preserved, with most of the mouthparts and epipharynx still attached, although the antennae are missing. Body size and sex remain unknown.

#### Etymology

The species name comes from the Latin word for “dragon,” reflecting its dragon-like head shape, unique among sisyphines.

### *Nesosisyphus vicinus* (Vinson, 1939)

Figs. 3a, 7d, 8a-e-i, 9a

*Sisyphus vicinus*: Vinson (1939): 36

*Nesosisyphus vicinus*: Vinson (1946): 92

*Nesosisyphus vicinus*: Vinson (1951): 93

#### Type locality

Maccabé forest (548 m), Mauritius

#### Size

Male and female: length: 3.4-4.4 mm; max. width: 2.0-2.7 mm.

#### Diagnosis

*N. vicinus* resembles *N. regnardi*. However, it can be easily recognized, in part, by a semi-circular incision between the median clypeal teeth, which is straight in *N. regnardi*; moreover, elytra are not unicolorous as in *N. regnardi* but variegated with yellow and brown interstriae.

#### Examined type material. Holotype

1 male, ADD-LOC, by photograph (MNHN). **Paratype:** Mauritius, Gorges Riv. Noire, 20.ii.1937, leg. J. Vinson (BMNH).

#### Examined non-type material

(30 ex. MZHF) Mauritius, Mt. Cocotte, 20°26’29.44”S 57°28’20.24”E, 15-17.ii.2021, 6 chicken dung baited traps, leg. S. Tarasov & M. Rossini; (34 ex. NHML, DMNS) Mauritius, Plaine Champagne, 20°25’S 57°25’E, vi.2004, chicken dung baited pitfall trap, leg. S. Motala.

#### Distribution

*N. vicinus* is by far the most widespread of the four extant Mauritian species. Historically, it was usually found in the tropical forests across the mountain peaks and slopes rising from the flat plateau areas in the southwest (Fig. 1b): Black River Gorges near Maccabé forest, Mt. Cocotte, Trou-aux-Cerfs (extinct crater); Grand Bassin peak, Bassin Blanc, Plaine Champagne. Previously recorded on a couple of isolated peaks in the center of the island: Piton du Milieu (590 m) and Trou-aux-Cerfs crater. Recent sampling efforts confirm the presence of *N. vicinus* in the south-western localities, but no further data are available from the central part of the island.

#### Remarks

Voucher specimen used for DNA extraction is deposited at MZHF under the following identification code: STL-1.

### *Nesosisyphus regnardi* (Alluaud, 1898)

Figs. 3b, 7e, 8b-f-l, 9b

*Sisyphus regnardi*: Alluaud C. (1898): 64

*Nesosisyphus regnardi*: Vinson (1946): 94

*Nesosisyphus regnardi*: Vinson (1951): 94

*Nesosisyphus regnardi*: Edmonds and Halffter (1978): 325, 328

#### Type locality

Mt. Le Pouce, Moka Range, Mauritius

#### Size

male and female: length: 3.3-4.9 mm; max. width: 1.8-2.7 mm.

#### Diagnosis

*N. regnardi* is morphologically similar to *N. rotundatus*. However, it can be distinguished from the latter by the combination of the following characters: the hind wings of *N. regnardi* are reduced to three quarters the length of the elytra while in *N. rotundatus* to very small rudiments about one-third of the length of the elytra 8f. Humeral and apical calli on elytra are still prominent unlike in *N. rotundatus* where they are absent. Further characters are given in the key.

#### Examined type material

(photograph of 1 Male, MNHN) Holotype: Mauritius, Mt. Le Pouce, 11.iii.1897, leg. G. Regnard.

#### Examined non-type material

(45 ex. MZHF) Mauritius, Mt. Le Pouce, 20°11’54.97”S 57°31’45.84”E, 12-15.ii.2021, human, chicken, monkey, tortoise dung baited trap, leg. S. Tarasov & M. Rossini; (127 ex. MZHF) Mauritius, Mt. Le Pouce, 20°11’54.99”S 57°31’45.84”E, 12-15.ii.2021, 13 chicken dung baited traps, leg. S. Tarasov & M. Rossini; (2 ex. MZHF) Mauritius, Mt. Le Pouce, 20°11’54.96”S 57°31’45.84”E, 21.ii.2021, 2 pitfall traps baited with carrion of land snails, leg. S. Tarasov & M. Rossini; (2 ex. MZHF) Mauritius, Mt. Le Pouce, 20°12’0.36”S 57°31’38.28”E, 12-25.ii.2021, hand collection, leg. S. Tarasov & M. Rossini; (6 ex. MZHF) Mauritius, Mt. Le Pouce, 20°12’6.86”S 57°31’28.92”E, 15-17.ii.2021, leaf litter sifting, leg. S. Tarasov & M. Rossini; (8 ex. NHML, DMNS) Mauritius, Mt. Le Pouce, 20°13’S 57°31’E, v.2004, pitfall trap baited with chicken dung, leg. S. Motala.

#### Distribution

*N. regnardi* occurs in the Moka range from Mt. Ory to Mt. Calebasses. In Mt. Le Pouce it occupies elevations above approximately 440 m (Fig. 1a-c). According to historical records, *N. regnardi* might be sympatric with *N. pygmaeus* and *N. rotundatus* in the middle of the southern slope of Mt. Ory.

#### Remarks

Voucher specimen used for DNA extraction is deposited at MZHF under the following identification code: STL-2.

### *Nesosisyphus rotundatus* Vinson, 1946

Figs. 3d, 7b, 8d-h-n, 9c

*Nesosisyphus rotundatus*: Vinson (1946): 95

*Nesosisyphus rotundatus*: Vinson (1951): 94

#### Type locality

S slope of Mt. Ory (441 m), Moka Range, Mauritius

#### Size

Male and female: length: 3.5-3.9 mm; max. width: 2.1-2.4 mm.

#### Diagnosis

Clypeus quadridentate, upcurved antero-laterally, lateral teeth short and blunt, medial teeth sharp and separated by straight-bottomed incision, Fig. 7b; surface of elytra smooth, without humeral or apical calli or shallow depressions like in the other species, Fig. 8d; hind wings reduced to very small rudiment about one-third length of elytra, Fig. 8h.

#### Examined type material

(1 ex. female NHML) Holotype: Mauritius, Mt. Ory (441 m), 18.xi.1944, leg. J. Vinson; (1 ex. female NHML) Paratype: Mauritius, Mt. Ory, i.1940, leg. J. Vinson.

#### Distribution

This species is highly localized on the southern slope of Mt. Ory, between 380 and 440 m. It was found where the ranges of *N. pygmaeus* and *N. regnardi* overlap.

#### Remarks

*N. rotundatus* is a comparatively rare species that was collected in only 6 specimens by J. Vinson during the early 1940s. This species has never been collected again, despite fieldwork and team efforts over the years on the island, including sampling at the type locality. Irreversible consequences combined with the uncontrolled habitat destruction in Mauritius led us to regard this species as possibly extinct. We used an archival protocol to extract and sequence the DNA from the holotype Lopes et al. (2023a). No secondary sexual characters have been detected since the examined material includes only females.

### *Nesosisyphus pygmaeus* Vinson, 1946

Figs. 3c, 7c-h, 8c-g-m, 9d

*Nesosisyphus pygmaeus*: Vinson (1946): 93

*Nesosisyphus pygmaeus*: Vinson (1951): 94

*Nesosisyphus pygmaeus*: Edmonds and Halffter (1978): 325

#### Type locality

S slope of Mt. Ory, Moka Range, Mauritius

#### Size

Male and female: length: 2.3-3.2 mm; max. width: 1.3-1.8 mm.

#### Diagnosis

Smallest known species of Sisyphini tribe. Clypeus bidentate, weakly upcurved anteriorly, incision between teeth semi-circular; humeral and apical calli prominent, the latter forming a tubercle protruding backward; hind wings and venation fully developed, three times lenght of elytra.

#### Examined non-type material

(1 ex. MZHF) Mauritius, Moka Range, S slope of Junction Peak, 20°12’44.36”S 57°29’51.58”E, 22.ii.2021, human dung baited trap, leg. S. Tarasov & M. Rossini; (1 ex. MZHF) Mauritius, Mare aux Songes, 20°26’48.44”S 57°41’52.56”E, 2007, subfossil in sediment sample, leg. N. Porch; (1 ex. NHML) Mauritius, Brise Fer Forest, 20°22’S 57°26’E, v.2004, pitfall trap baited with chicken dung, leg. S. Motala.

#### Distribution

Despite being fully winged, *N. pygmaeus* was known only from the restricted area of Mt. Ory until further samplings in 2003, led to the discovery of a new population in a new locality approximately 24 kilometers away from Mt. Ory in Brise Fer Forest, 20°22’S 57°26’E (Motala and Krell, 2007).

On the S slope of Mt. Ory the population of *N. pygmaeus* lives together with *N. rotundatus* (old records) and *N. regnardi*.

#### Remarks

Among the subfossils collected in Mare aux Songes, a small pronotum of *Nesosisyphus* sp. was found, see Fig. 7h. Comparing the pronota of all species, *N. pygmaeus* appears to be closest in punctuation, shape and size (length and width of pronotum). As *N. pygmaeus*, with a body size of 2 to 3 mm, is the smallest species within the genus, we assume that the pronotum found belongs to this species based on the matching size and morphological characters.

Voucher specimen used for DNA extraction is deposited at MZHF under the following identification code: STL-3.

## DISCUSSION

### Systematic and taxonomy of *Nesosisyphus*

In the present paper, we focused the phylogenetic analysis on clarifying the systematic position of *Nesosisyphus* by incorporating species with African, Palearctic, and Oriental distribution. The three loci-based phylogenetic results (Fig. 5) reveal that *Nesosisyphus* forms a strongly supported monophyletic group, yet it is nested within the genus *Sisyphus*, which also includes African and Palaearctic species. This finding highlights potential taxonomic changes. The nesting of *Nesosisyphus* within *Sisyphus* suggests that the latter, as currently defined, may be paraphyletic. Resolving this taxonomic inconsistency requires either reclassification of *Nesosisyphus* as a subgenus within *Sisyphus* or redefining genus boundaries. The UCE-based phylogeny (Fig. 4) further provides a markedly different perspective on the relationships within the tribe Sisyphini, aligning more closely with the scenario of *Nesosisyphus* as an independent lineage. In this analysis, Sisyphini is recovered as a sister clade to the tribe Epirinini, with a distinct split observed within Sisyphini. One group, clade A (BS = 100%), comprises exclusively all *Nesosisyphus* species. This finding contrasts with the earlier concatenated phylogeny, which nested *Nesosisyphus* within *Sisyphus*. The recovery of *Nesosisyphus* as an entirely distinct monophyletic clade from *Sisyphus* provides strong support for its evolutionary independence, suggesting a biogeographical scenario where an ancestral population colonized Mauritius and subsequently evolved in complete isolation. The robustness of UCE-based phylogenies has been demonstrated in several studies. For instance, (Gilbert et al., 2015) highlighted the effectiveness of UCEs in resolving complex phylogenetic relationships due to their cost-effectiveness and high resolution. Similarly, (Blaimer et al., 2015) and (Branstetter et al., 2017) demonstrated the advantages of UCEs in resolving both ancient and recent divergences in formicine ants, yielding increased node support and clearer taxonomic insights. Our UCE-based results, based on 3,000 loci target set, provide therefore a stronger and more reliable phylogenetic framework compared to singlegene phylogenies. To present, clade A represent a lineage exclusive to *Nesosisyphus*, solidifying its status as a separate genus and highlight a significant divergence within Sisyphini. Phylogenomic approaches that integrate data from multiple loci, such as UCE analyses, tend to provide a more comprehensive perspective on species evolution, as they are less susceptible to the biases inherent in single-gene markers. If incomplete lineage sorting is not acting massively across genomes, broader genomic sampling generally leads to a more accurate representation of species relationships. Given the robustness and reliability of UCE-based phylogenies, we therefore base our main conclusions on these results, supporting *Nesosisyphus* as an independent lineage. Nevertheless, the discrepancies observed across different genetic markers highlight the challenges of phylogenetic reconstruction. Although single-gene analyses contribute with valuable information, phylogenomic approaches that integrate multiple genetic markers offer a more holistic perspective on evolutionary relationships. Differences in phylogenetic results could be attributed to various factors, including dataset composition and methodological approaches. Indeed, further investigations are required: molecular dating and comprehensive dataset including Neotropical species of Sisyphini are planned for future analysis.

While divergence time estimation was not the focus due to data limitations, the longer branches observed on *Nesosisyphus* (Fig. 4, 5) may indicate a basally originating lineage that diverged relatively far in the past, probably from African, where most species are found. Nevertheless, this question remain for the ongoing work and will be better solved in subsequent phylogenetic analysis.

A series of morphological features were listed to justify *Nesosisyphus* as a separate genus (Vinson, 1946). We examined and evaluated those characters and, based on the UCEs molecular results (Fig. 4), we support their validity in distinguishing *Nesosisyphus* from all the other species.

The head of the new subfossil species *N. draco* n. sp. (Fig. 7f-g), was found among a surprising quantity of insect remains, accumulated within a small time window between 4400 and 3900 cal. yr BP (Rijsdijk et al., 2011). The excellent conditions and the importance of head morphology in species identification within the Sisyphini, led us to describe it as a new species. Unfortunately, since the species is known from a subfossil and only from the lowlands, we must conclude that is likely extinct.

A subfossil pronotum, found in the same fossil bed, appears to have a morphology closely related to the one of *N. pygmaeus*, based on the matching size and pronotal punctuation Fig. 7h. On the other hand, the anterior margin of the pronotum could match with the head of *N. draco* n. sp. However, since the insect subfossils are found in disarticulated body parts, the prothorax of *N. draco* n. sp. remains unknown, and our consideration is based on the available data from extant species.

### The evolution of *Nesosisyphus* in Mauritius

The timing of *Nesosisyphus* arrival to Mauritius was not within the scope of this paper, as it requires comprehensive dating analyses and future investigation. In this paper we instead discuss the events following *Nesosisyphus* arrival, which obviously occurred after the emergence of Mauritius around 10 Mya. *Nesosisyphus* populations on the island exhibit generalist feeding behavior, attracted to a wide range of dung sources without strong preferences. Previous studies suggest reptiles and fowl as primary dung sources to maintain populations Lopes et al. (2023b). Precisely, the species of *Nesosisyphus* might have shaped their diet as their habitat became restricted and key-species of vertebrates got extinct. Vinson (1947) formerly hypothesized an original diet based on the excrement of the Dodo (*Raphus cucullatus*) and the giant land tortoises (*Cylindraspis* spp.), although our evidence indicates that dung from current large tortoises from the Seychelles does not attract *Nesosisyphus*, possibly due to the extinction of specialized species. This hypothesis was raised mainly due to the lack of large native mammals in Mauritius (Cambefort and Hanski, 1991). Species that evolved in certain environments (e.g. islands, rainforest) with absence of indigenous large mammals, possibly adapted to different food resources (generalist feeders) as a response to the lack of large mammal dung (Ebert et al., 2019; Langton-Myers, 2022; Cupello, 2023). Based on recent studies on dung beetle feeding habits in islands without large native mammals, an original diet on large bird droppings and reptile dung is feasible (Jones et al., 2012; Stavert et al., 2014; Lopes et al., 2023b). In Mauritius island, when human colonization started, many vertebrate species were exterminated, including the Dodo and the giant land tortoises. Therefore, present-day feeding habits of *Nesosisyphus* species might be quite different from how they used to be in origin. For example, we collected a relatively large number of specimens of *N. regnardi* by using chicken droppings, and to less extent with the carrion of the invasive snail *Lissachatina fulica* (Lopes et al., 2023b) *and human dung. From the bait experiment, several specimens of N. pygmaeus* were attracted mainly to avian fauna (rotting chicken meat and chicken droppings), bat and crocodile feces (endemic bat + avian fauna = 78% of captures), and to a lesser extent to monkey (*Macacus cynomolgus*, an Indian species introduced in Mauritius in the early 16th century) feces and snail carcasses. None specimens were collected on giant tortoise *Aldabrachelys gigantea* (the endemic species to the Seychelles) dung-baited pitfall traps. N.P. observed living adults of *N. vicinus* feeding on monkey’s excreta (Porch, 2016). Also Vinson (1939) reported observations of *Nesosisyphus* adults feeding on monkey’s excreta and the carrion of the endemic Mauritian mollusc *Pachystyla bicolor* (Vinson, 1951), *and collected several specimens using fowl dung as bait for pitfall traps. Given the Nesosisyphus* preferences for avian dung and their ability to use local resources in the absence of large mammals, it might be likely that the largest Mauritian bird droppings (for instance, the Dodo bird) have served as food resource.

While most of the known *Nesosisyphus* species have survived to human impact by shaping their feeding habits and ranges, subfossil evidence indicates that the dung beetle diversity in Mauritius was once richer in terms of species number. Indeed, the paleontological survey conducted in MAS site produced a large number of arthropod subfossils, revealing at least five new subfossil species of Scarabaeinae (N.P. unpubl. data), including the newly described Sisyphine. It is noteworthy that mass mortality events of the native fauna, such as at MAS, are attributed to natural megadrought that occurred in Mauritius about 4000 yrs ago, and did not result in the extinction of any of the species involved, but rather in the deaths of many individuals locally (Rijsdijk et al., 2011). Many of the species involved in this natural event became extinct soon after the arrival of humans as a result of hunting, invasive species and habitat loss (for instance, Fig. 6A shows the profound changes in the indigenous forest cover during human expansion). Radiocarbon dating (Rijsdijk et al., 2015) and subfossil evidence (Rijsdijk et al., 2011) provide further support for this hypothesis. Moreover, when there are continuous records, such as from Rodrigues and many islands in the Pacific, the timing of extinctions always correlates with human arrival (Martin and Klein, 1984; James et al., 1987; Burney et al., 2004; Steel, 2015; Matthews and Triantis, 2021). In Mauritius we haven’t managed to find a continuous sequence rich in insects, so it is impossible to be definitive regarding the specific timing of extinction on those dung beetles from a sample that dates to one small part of the past. We rather use evidence from other taxa (the vertebrates) and other localities (such as Asia, East Africa and Madagascar) to suggest the likely scenario, which is extinction and loss of lowland biota linked to human disturbance. It is likely that a similar situation affected the dung beetle fauna in the other two Mascarene islands of La Réunion (Rossini et al., 2021) and Rodrigues (Nick Porch unpubl. data).

Although the insect remains themselves have not been dated, their occurrence in sediments that have been dated to this period suggest they are likely to be the same age. As such they provide unique evidence of the nature of the Mauritian lowland insect fauna prior to the destruction of the lowland forests (Hammond et al., 2015).

Our results based on niche modeling suggest that the distribution pattern of *Nesosisyphus* changed in a significantly short time in the past 400 years. This group of dung beetles used to be widely distributed across the island (Fig. 6B), even on the uplands of the eastern side, where S.T. and M.R. conducted several samplings, without success. The model shows also different ecological niches for *N. vicinus* and *N. regnardi*, which may have partially overlapped in the past. Present-day populations are instead restricted to the best preserved areas of the island, with different species occupying different mountain blocks, primarily located on a few mountains in the western side, and confined to disturbed patches of tropical forest, see Fig. 2. It is unlikely that such a recent isolation was caused by long-term natural events. Vertebrate extinctions and forest contraction driven by human expansion deeply modified the habitat with possible consequences on the relationships between the species. It is known that species extinction and relict populations among dung beetles can be the result of a series of cascading negative impacts across various species groups, given that dung beetles typically rely on excrement as food resource (Culot et al., 2013). Therefore, present microendemisms might have been promoted by interspecific competition at a local scale due to the lack of resources, and are most likely the remnant of local extinctions. Additionally, this process may have been fostered by the flightlessness of some species. For instance, the case of the flightless *N. rotundatus* is emblematic. Even the subfossil record of *N. pygmaeus* in MAS site indicates a wider distribution in the past, and might be another warning signal of the growing threat that these dung beetles are currently facing. Additionally, the lack of records for *Nesosisyphus* in Eastern Mauritius, where the habitat shows suitable niches, manifest a possible ongoing decline.

At least one subfossil species of *Nesosisyphus* is extinct, and another one is most likely extinct. Our predictive species distribution model and subfossils findings prove how the dung beetle fauna used to be spread in Mauritius, in both lowland and upland, and how all declined precipitously, coincident with the human arrival.

## CONCLUSIONS

In the present paper, we used molecular analysis to assess the systematic position of the genus *Nesosisyphus* endemic to Mauritius, with the description of a fifth new species from a subfossil.

The phylogenetic analysis based on UCEs recovered *Nesosisyphus* as a distinct lineage, separated from other Sisyphini lineages, whereas the three-loci phylogeny places *Nesosisyphus* as a monophyletic group within the genus *Sisyphus*. In cases where different analyses yield conflicting evolutionary interpretations, it is essential to prioritize the method that offers greater reliability. UCE-based phylogenies have been demonstrated to be more robust in multiple studies. Phylogenomic approaches that integrate data from multiple loci, such as UCE analyses, provide a more comprehensive perspective on species evolution, as they are less prone to the biases inherent in single-gene markers. Therefore, we base our conclusions on the UCEs results, solidifying its status as a separate genus and highlights a significant divergence within Sisyphini.

To assure accuracy in divergence time estimation and to assess the biogeographic origin, we propose the inclusion of Neotropical Sisyphini in future phylogenetic analyses.

The taxonomic investigation agrees with Vinson’s previous species concepts, and no other new extant species were discovered on the island. However, the examination of subfossils revealed a new species, *Nesosisyphus draco* sp. n., along with another currently existing species, likely belonging to *N. pygmaeus*. The discovery of Sisyphine subfossil sclerites underscores the long-standing presence and evolutionary significance of this group on the island. This finding further highlights the importance of integrating paleontological data with contemporary surveys to reconstruct historical baselines and assess biodiversity changes over time. We also identified that the flightless *N. rotundatus*, collected only in 1940s, may be extinct at present due to habitat loss. Also the newly described species, *N. draco* n. sp., is regarded here as an extinct species since is known just from a 4000 yrs old subfossil sclerite. These findings suggest that multiple extinction events have influenced the diversity and distribution of *Nesosisyphus* after the human colonization. Indeed, Mauritian Sisyphines are currently found only in a few isolated patches of mountainous forest throughout the central plateau, exactly where the best preserved habitats occur.

These findings emphasize the importance of conserving *Nesosisyphus* and its habitats. Future efforts should focus on further studies to assess the species’ current status (IUCN extinction risk assessment) and ecological needs.

We performed a niche modeling analysis to estimate the potential ecological niches of *Nesosisyphus* species and conducted feeding experiments in the field to discuss how the impact of humans might affected negatively their distribution and diet preferences.

## DATA AVAILABILITY STATEMENT

Raw sequenced data are available in GenBank database under the accession numbers: PP234995, PP234996, PP234997 (COI gene sequences); PP239338, PP239339, PP239340 (16S gene sequences); PP257798, PP257799, PP257800 (28S gene sequences).

## AUTHOR CONTRIBUTIONS

FLos contributed to the analysis and interpretation of data, visualization of data, and was in charge of writing the original manuscript. FLop contributed to the analysis of data and edited the original draft. MR contributed to the acquisition of data, and revised the original manuscript. NP, FTK, and SM contributed to the acquisition of data, funding acquisition, and edited the original manuscript. GMD revised/edited the original manuscript. ST designed the study, contributed to the acquisition of data and funding, was responsible for the analysis and interpretation of data and implemented the original draft adding significant contributions. All the authors have read and approved the final version of the article.

## ACKNOWLEDGMENTS

The authors want to thank the Mauritian National Parks and Conservation Service (NPCS) for providing permits to collect in Mauritius; Vincent Florens (University of Mauritius) and Owen Griffiths (The Australian Museum) for logistic support, continuous help during our 2021 expedition to Mauritius and sampling efforts in 2022; Saoud Motala’s fieldwork was supported by Vikash Tatayah, Jean Claude Sevathian (Mauritian Wildlife Foundation) and by the National Parks and Conservation Service; we would like to thank Kenneth Rijsdijk (University of Amsterdam) for providing sediment samples from the MAS excavation to Nick Porch for insect analysis; Max Barclay (NHML) for the loan of type material of *N. rotundatus*; Olivier Montreuil (Muséum National d’Histoire Naturelle, Paris) for providing access to the type specimens of *N. regnardi* and *N. vicinus*; Hennie de Klerk and Conrad P.D.T. Gillett for the live photographs of “Other Sisyphini” used in Fig. 4; Pedro Cardoso for useful insights on the niche modeling; Dmitry Zastrozhnov for guiding us in processing the DTM elevation model of Mauritius used in Fig. 2.

## SUPPLEMENTARY MATERIAL

Additional information about this study are freely available online at: https://osf.io/37rmj/.

